# Attention Networks and the Intrinsic Network Structure of the Human Brain

**DOI:** 10.1101/2021.06.01.446566

**Authors:** Sebastian Markett, David Nothdurfter, Antonia Focsa, Martin Reuter, Philippe Jawinski

## Abstract

Attention network theory states that attention is not a unified construct but consists of three independent systems that are supported by separable distributed networks: an alerting network to deploy attentional resources in anticipation of upcoming events, an orienting network to direct attention to a cued location, and a control network to select relevant information at the expense of concurrently available information. Ample behavioral and neuroimaging evidence supports the dissociation of the three attention domains. The strong assumption that each attentional system is realized through a separable network, however, raises the question how these networks relate to the intrinsic network structure of the brain.

Our understanding of brain networks has advanced majorly in the past years due to the increasing focus on brain connectivity. It is well established that the brain is intrinsically organized into several large-scale networks whose modular structure persists across task states. Existing proposals on how the presumed attention networks relate to intrinsic networks rely mostly on anecdotal and partly contradictory arguments. We addressed this issue by mapping different attention networks with highest spatial precision at the level of cifti-grayordinates. Resulting group maps were compared to the group-level topology of 23 intrinsic networks which we reconstructed from the same participants’ resting state fMRI data. We found that all attention domains recruited multiple and partly overlapping intrinsic networks and converged in the dorsal fronto-parietal and midcingulo-insular network. While we observed a preference of each attentional domain for its own set of intrinsic networks, implicated networks did not match well to those proposed in the literature. Our results indicate a necessary refinement of the attention network theory.

## 1. Introduction

In order to react adequately and to act purposefully in a dynamic and ever-changing environment, the brain needs to prioritize information processing, e.g. by anticipating when and where sensory information will appear, or by selecting more relevant over less relevant information. Attention refers to the cognitive function that guides the prioritization and selection of some at the expense of other information (Cowan, 1999; Posner & Fan, 2008). Converging evidence from single cell recordings, electrophysiology, and neuroimaging suggests that ongoing neural information processing is enhanced in a highly specific and targeted way when attention is shifted towards a certain location in the visual field (Brefczynski & DeYoe, 1999; Heinze et al., 1994; Kastner et al., 1999; Luck et al., 1997; Müller et al., 2003) or towards task-relevant stimulus features (Egner & Hirsch, 2005; O’Craven et al., 1997; Rees et al., 1997). While the effects of attention become apparent in increased firing rates and BOLD activity in sensory areas which process the currently attended information, the recruitment and control of attention signals is realized by neural systems that additionally include areas upstream on the cortical processing hierarchy (Posner & Petersen, 1990, Posner & Dehaene, 1994). Attention network theory assumes three largely independent systems that realize one out of three different types of attention: the alerting system initiates a state of increased arousal in direct anticipation of upcoming stimuli, the orienting system shifts the attentional focus to locations in space, and the control system selects and amplifies relevant information when distracting or task-incompatible information is present (Posner & Petersen, 1990). The three systems are thought to dissociate neuroanatomically into independent ‘attention networks’ (Posner & Rothbart, 2007). Evidence for the relative independence of the three attention systems comes from research with the attention network test (Fan et al., 2002). The attention network test (ANT) is a reaction time task that combines the flanker task (Eriksen & Eriksen, 1974) to study the attentional selection of relevant information at the expense of irrelevant distractors with the Posner cueing task (Posner, 1980) where briefly presented cues carry information when and where an upcoming target stimulus will appear. Behavioral indices of the efficiency of alerting, orienting, and control are uncorrelated (Fan et al., 2002) which is interpreted as an indication of independent systems. Moreover, genetic work points towards different genetic contributions and underlying susceptibility variants for the attention systems (Fan et al., 2001; Fossella et al., 2002; Reuter et al., 2007). Furthermore, neuroimaging work with the ANT has revealed non-overlapping activation patterns for task contrasts that probe alerting, orienting, and attentional control (Fan et al., 2005), adding further evidence for dissociable and presumably independent systems. At the measurement level, attention networks are often equated with the activation patterns elicited by the ANT. The focus on task activations and the negligence of functional connectivity for network definition raises the question how the ANT-attention networks relate to the intrinsic network structure of the human brain.

Our understanding of brain networks has advanced majorly in the past years due to the increasing focus on brain connectivity. Several large scale networks that delineate along functional boundaries of the brain have been identified in spontaneous intrinsic BOLD fluctuations in the task-free resting state (Fox & Raichle, 2007; Smith et al., 2013; van den Heuvel & Hulshoff Pol, 2010). Importantly, the intrinsic network architecture persists into task states and matches the topology of task-evoked activations (Cole et al., 2014; Gordon et al., 2012; Nickerson, 2018; Smith et al., 2009). If the three attention systems were actually independent networks, we would assume that each system activates a distinct or distinct group of intrinsic connectivity network (ICN). This has also been suggested previously, for instance, that the three attention networks segregate within an ‘extended fronto-parietal network’ (Xuan et al., 2016), that the orienting network corresponds to a dorsal and a ventral fronto-parietal network and the attention control network to a distinct fronto-parietal and an insular-opercular network (Petersen & Posner, 2012). The fronto-parietal and insular-opercular network have also been discussed regarding their role in alerting (Sadaghiani & D’Esposito, 2014). Some of these previous propositions, however, rely only on anecdotal arguments and appear in conflict with each other. At present, it is unclear how the idea of three separable and independent attention networks as activated by the ANT is reflected in the overall network structure of the brain. We designed the current study to directly probe the spatial correspondence between ICN and the three attention networks. We first recorded resting-state fMRI data in order to delineate ICN and second recorded task fMRI data from the same participants with the most recent version of the ANT (the revised ANT, (Fan et al., 2009; Xuan et al., 2016). We made use of recent developments by the Human Connectome Project to achieve high spatial precision through multimodal surface matching (Robinson et al., 2018) and minimal spatial smoothing (Glasser et al., 2016).

## 2. Methods

### 2.1 Participants

We recruited N = 86 healthy young adults (age: M = 26.17 years, SD = 5.41 years; n = 40 females, n = 46 males) through flyer advertisements on campus, mailing lists, and announcement in undergraduate psychology classes. Participants were screened during a telephone interview to meet the following inclusion criteria: Native-level proficiency in German, right-handedness, and age between 18 and 35 years. We targeted an equal amount of male and female participants. Participants were excluded when they indicated past or present psychiatric or neurological illness, psychotropic substance use in the past six months, or any contraindication to MRI. Participants reported to have normal or corrected-to-normal vision during the experiment. Informed written consent was obtained prior to enrollment in the study. Participants were remunerated with the usual rate of 10 EUR/hour (i.e. 25 EUR for the entire study) or its equivalence in course credit, if desired by the participant. The study protocol was in accordance with the Declaration of Helsinki and approved by the ethics committee of the University Hospital Bonn.

### 2.2 Attentional Network Test

We adapted the Attentional Network Test in its revised form (Xuan et al., 2016). The ANT-R combines a spatial cueing with a flanker task and is the standard protocol to activate different attentional systems in the brain. We administered a total of 288 trials in four runs of 72 trials each. A typical trial sequence is shown in figure 1. The ANT-R follows a 4 x 2 design with the factors cueing condition (no cue, double cue, valid spatial cue, invalid spatial cue) and target (congruent flanker, incongruent flanker). Activation maps and behavioral indices for the attention networks were computed by contrasting different cue and target conditions as described below (see task analysis and behavioral analysis). Each run lasted for 420 seconds, leading to a total time of around 30 minutes for the whole experiment.

**Figure 1:**
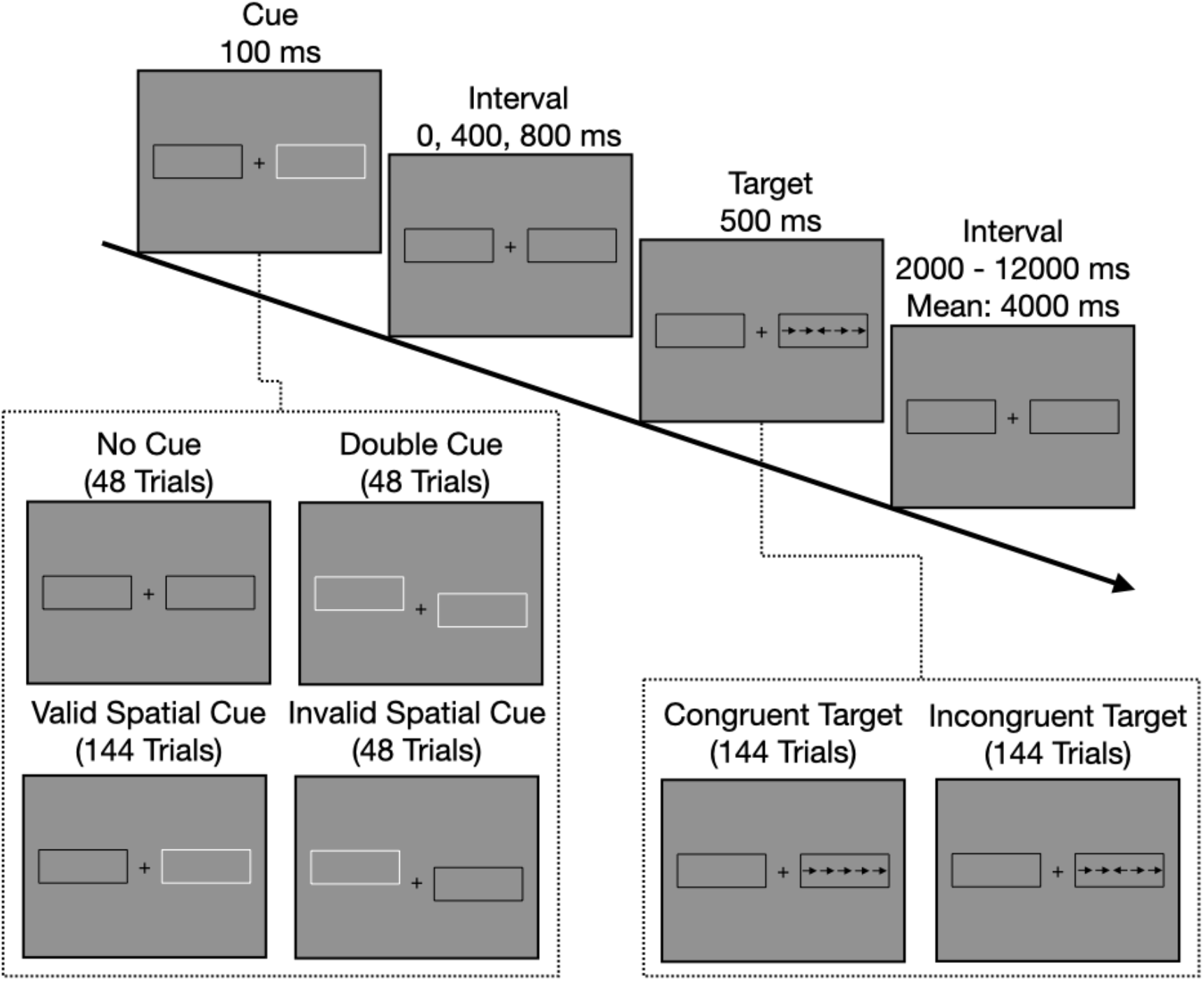
Schematic overview over stimuli and stimulus timing in a typical trial sequence. Each trial started with a 100 ms presentation of either no cue, a double cue, or a spatial cue. After a cue-target interval of 0, 400, or 800 ms, five arrows were flashed for 500 ms as target stimulus. Participants indicated via button presss whether the central arrow pointed to the left or to the right. Flanking arrows were either congruent or incongruent (half of the trials each). Target offset and onset of the next cue were spaced by a jittered interval (mean interval across trials: 4000 ms, range: 2000 to 12000 ms). Targets appeared either at the cued position (valid spatial cues) or at uncued position (invalid spatial cue). A total of 288 trails was presented in four runs.

Throughout each run a fixation cross was presented in the middle of the screen, surrounded by a rectangle on its left and right side (the rectangles subtended 4.69° of visual angle to both sides). The fixation cross and the rectangles remained visible during the whole run. In every trial, arrows were presented in one of the rectangles: An arrow in the center (target) was surrounded by two arrows each on the left and the right side (flankers). Each arrow subtended 0.58° of visual angle and the distance between arrows was 0.06° of visual angle. The arrows pointed either to the left or to the right and the five arrows could either be congruent (i. e. the target arrow pointed to the same direction as the surrounding flankers) or incongruent (i.e. the target arrow pointed to the opposite direction as the flankers). Participants were instructed to select as fast and accurately as possible the direction of the middle arrow by either pressing a button in the left or the right hand. In some trials, a cue was presented before the flankers appeared via brightening of one or both of the rectangles. As a spatial cue, only one of the rectangles flashed, while a brightening of both rectangles (double cue) served as a temporal cue. Spatial cues could either be valid, i.e. the arrows were presented in the rectangle that brightened, or invalid, i.e. the arrows were presented in the opposite rectangle. A short interval was implemented between cue and flanker presentation.

Each trial consisted of three phases: a cue phase (100 ms), a short interval (0, 400 or 800 ms, equally distributed) and a target phase (500 ms). The different conditions were spread across two blocks consisting of 144 trials: Of the 144 trials, one sixth, i.e. 24, were no cue, double cue and invalid spatial cue trials, respectively. The other half, i.e. 72 trials, consisted of valid spatial cues. Each cue type was followed by each other cue type equally often (Fan, 2009). The 24 combinations of interval between cue and target phase, flanker type (congruent or incongruent) and target location (left or right rectangle) were randomized for each cue condition. The interval between offset of target and onset of the next trial was distributed systematically between 2000 and 12,000 ms with a mean of around 4000 ms (for details see Fan et al., 2009). While the target was only presented for 500 ms, participants had additional 1200 ms to press the button after offset of the target, leading to a total time frame of 1700 ms to respond.

The experiment was programed with Presentation software version 20.1 (Neurobehavioral Systems, Inc., Albany CA) and presented via a projector in the MR scanner. The projection screen had a resolution of 1024×768px (24×18cm) and the distance between screen and participants’ eyes was around 62 cm. Participants took part in a short training block consisting of 10 trials outside of the MR scanner to get familiar with the setup.

### 2.3 Image Acquisition

All MR images were acquired in a single session on a Siemens 3T Prisma equipped with a 32 channel head coil at the Berlin Center for Advanced Neuroimaging between March and December 2019. We adopted MR sequences from the HCP-Lifespan project (Harms et al., 2018). The following protocols were acquired in a fixed order: 1) T1-weighted structural (Multiecho MPRAGE, voxel size 0.8mm isotropic, time to repeat TR=2.4s, time to echo TE=22ms, flip angle 8°) 2) T2-weighted structural (SPACE, voxel size 0.8mm isotropic, TR=3.2s, TE=563ms, flip angle 120°), 3) BOLD rfMRI (multiband echoplanar, 72 slices, 805 volumes, TR=800ms, voxel size 2mm isotropic, TE=37ms, flip angle 52°, A-P encoding direction) including two spin echo fieldmaps (A-P and P-A encoding), 4) tfMRI in four runs with run-specific spin echo fieldmaps and the same pulse sequence as for rfMRI 4) Diffusion-weighted images (DWI). DWI data will not be part of the present report. A reference image without multiband acceleration was acquired for each functional run.

### 2.4 Preprocessing

We adapted the HCP minimal preprocessing pipelines (github.com/Washington-University/HCPpipelines) for structural and functional preprocessing (Glasser et al., 2013). If not stated otherwise, we used version 4.1 of the pipelines, Freesurfer 6.0.0, and FSL 6.0.1 under Linux Debian 10. Structural images (T1 and T2) were corrected for gradient distortions, aligned, brain extracted, bias field corrected, and registered to MNI space using non-linear transformation. Structural images where then further processed with HCP’s Freesurfer pipeline with improved brain extraction, alignment, and adjustment of the white matter surface. The Freesurfer output was converted to Nifti and Gifti files and used to create a brain mask for all further analyses. Cortical surfaces were then registered to template space based on cortical folding (MSMsulc, Robinson et al., 2018) and downsampled to the 32k_LR surface space. All functional data (rfMRI and task fMRI) and the corresponding field maps were processed with the fMRIVolume pipeline which included correction for gradient distortions, motion, EPI image distortions, co-registration with the T1 structural image, and normalization to MNI volumetric space. All transformations were applied in one step. Functional data were then intensity normalized to their global 4D mean and masked. The resulting volume timeseries were further processed with the fMRISurface pipeline to create individual CIFTI dense timeseries grayordinate files by resampling subcortical gray matter voxels to standard subcortical parcels and by partial-volume-weighted and cortical-ribbon-constrained-mapping of cortical gray matter voxels onto standard surface vertices. In this step, we applied light volume- and surface-based smoothing with a Gaussian filter with 2 mm full width at half maximum.

Resting state timeseries were processed further to remove artifacts. Each participants’ volumetric rfMRI timeseries were first run through FSL’s Multivariate Exploratory Linear Optimized Decomposition into Independent Components (MELODIC) tool (ve3.15) and the processed using FSL Fix (v1.06.15). We used a classifier that had been trained on the HCP young adult sample as distributed with FIX. Automatic component classification worked excellent despite small differences in acquisition parameters between our data and the training data. Manual inspection indicated that no component had to be re-labeled. Artifactual components were regressed out together with the six head motion parameters and their first temporal derivatives. The cleaned rfMRI time series were then converted to grayordinates as described above. The FIX pipeline was run on a twelve-core Mac Pro (High Sierra 10.12.6) machine using R (v3.3.3), Matlab (v2018b), and HCPpipelines (v4.2.1). Relevant R-packages were used in the respective version mentioned in the FIX documentation and re-compiled when needed.

### 2.5 Independent Component Analysis

We identified ICN at the group level by running a group ICA in MELODIC after concatenating all participants’ FIX-cleaned dense timeseries grayordinate files in time and reducing the data matrix into a 1609 dimensional subspace. We requested 27 components, after estimating the ICA’s dimensionality in Matlab using HCP code (icaDim.m). Four artifactual noise components were identified through visual inspection and the remaining 23 components were kept for further analyses.

We labeled these ICN based on a) the similarity of associated time courses obtained through hierarchical clustering of the IC time courses based on the time courses’ full correlation network matrices using the Ward method in FSLnets (v0.6.3) (see figure 3), b) pairwise comparison of each thresholded and binarized (|z| > 3) ICN with published atlases (the Cole-Anticevic cortical and subcortical partition, the Yeo 7 and 17 cortical networks, and the Power partition, see supplementary table T1), and c) visual inspection and comparison with a detailed map of cortical areas (see supplementary figure S1).

### 2.6 Task Analyses

First level analyses were performed in SPM12 (www.fil.ion.ucl.ac.uk/) using a general linear model. Surface images were converted to “fake-volumetric” nifti-images using wb_command. Condition specific regressors were created by convolving a train of delta functions with SPM’s canonical hemodynamic response function. Separate regressors reflected the onsets of the following events: double cues, no cues, valid cues, invalid cues, and congruent as well as incongruent targets following different cues. This resulted in 12 regressors. We added one additional regressor with the onsets of error trials, twelve regressors with the six head motion parameters and their temporal derivatives, and one constant per run. The final design matrix contained (12 + 13 + 1) * 4 columns.

Linear weighted contrasts were computed on the estimated beta images to derive the attention network maps (see Xuan et al., 2016): Alerting network: Double cue condition vs. implicit baseline. Control network: all incongruent targets minus all congruent targets (across cue conditions). The orienting network was operationalized via the Validity effect (invalid cue minus valid cue), which is a combination of disengaging attention from an invalid location (invalid cue minus double cue, the Disengaging effect) and moving and engaging the attentional focus to a validly cued location (valid cue minus double cue, the Moving + Engaging effect). Individual contrast images were back-converted to cifti-files and then passed on to second level group analysis.

We used the Sandwich Estimator (SwE) Toolbox for SPM12 (Guillaume et al., 2014) for group-level analyses of individual contrast images to assess activation of attentional systems across all participants. SwE’s main application is longitudinal and repeated measures neuroimaging data, but SwE is also suitable for more simple designs like ours. We used the modified SwE procedure with a small sample size correction (type c) and a wild bootstrapping procedure with 999 bootstraps. The family-wise error was corrected at the cluster level (p < .05) with a cluster-forming threshold of p < .01. A cluster forming threshold of p <.01 is rather liberal, however, the ANT is regarded a well-established paradigm and a similar threshold has been used in previous reports (e.g. Xuan et al., 2016).

### 2.7 Spatial Regression

Our main question focuses on the spatial relationship between intrinsic ICN and the different attentional “networks” as activated by the ANT-R. We used a multiple spatial regression approach (Gordon et al., 2012) to predict group-level ANT-activation maps from the 23 group-level ICN. Separate models were estimated for each activation map. Unthresholded activation maps (z-images) were reshaped to column vectors that included all cortical vertices and subcortical voxels. These vectors served as criterion in the regression analyses. On the predictor level, all 23 non-artifactual unthresholded IC maps were reshaped into a grayordinate * component matrix. Ordinary least square regressions were fitted using Matlab’s fitlm function. Possible confounds due to collinearity were ruled out by inspecting condition indices and variance decomposition proportions from the predictor matrix. Effect sizes for individual ICN were calculated as partial regression coefficients by residualizing each ICN from all remaining ICN, fitting a linear regression model, and obtaining the adjusted R2.

### 2.8 Behavioral Analysis

We analyzed reaction times and error rates to compute behavioral indices of the different attention networks. For reaction time analyses, all error trials and responses outside a response window of 1700 ms after target onset were excluded (see Xuan et al., 2016). Behavioral indices were computed as differences between experimental conditions: Alerting: no cue minus double cue, Orienting: Invalid Cue minus Double Cue (Disengaging), Double Cue minus Valid Cue (Moving + Engaging), and Invalid Cue minus Valid Cue (Validity Effect), and Control: Incongruent minus congruent target. We calculated five (attention contrasts) * two (reaction times, error rates) one-sample t-tests to test whether the behavioral index differed significantly from zero.

### 2.9 Final sample

We had to exclude two participants because of incidental findings. The final rfMRI sample included N = 84 (mean age M =26.34, SD =5.35, n = 47 female, n = 37 male). Six participants were excluded from the task analysis for committing an excessive number of errors in the ANT (n = 4), large artifacts in the task fMRI data (n=1), and incomplete task fMRI data (n = 1). The final tfMRI sample included N = 78 subjects (mean age M =26.19, SD =5.34, n = 44 female, n = 34 male).

### 2.10 Code and data availability

All preprocessing code for structural and functional preprocessing can be obtained from https://github.com/Washington-University/HCPpipelines. Group-level activation maps and ICN maps (thresholded and unthresholded) and Matlab code to match ICN to templates, run the spatial regression analysis and subsequent analyses will be made available on the Open Science Framework upon publication.

## 3. Results

### 3.1 Behavioral Results

Mean differences in reaction times and committed errors are presented in Figure 2. As expected, the presence of temporal and valid spatial cues led to faster reaction times while invalid cues and incongruent flankers led to slower responses (all p<.001). The same pattern was also visible in the error rates, except for the alerting contrast where the difference was not statistically significant (p = .47, all other p < .05).

**Figure 2:**
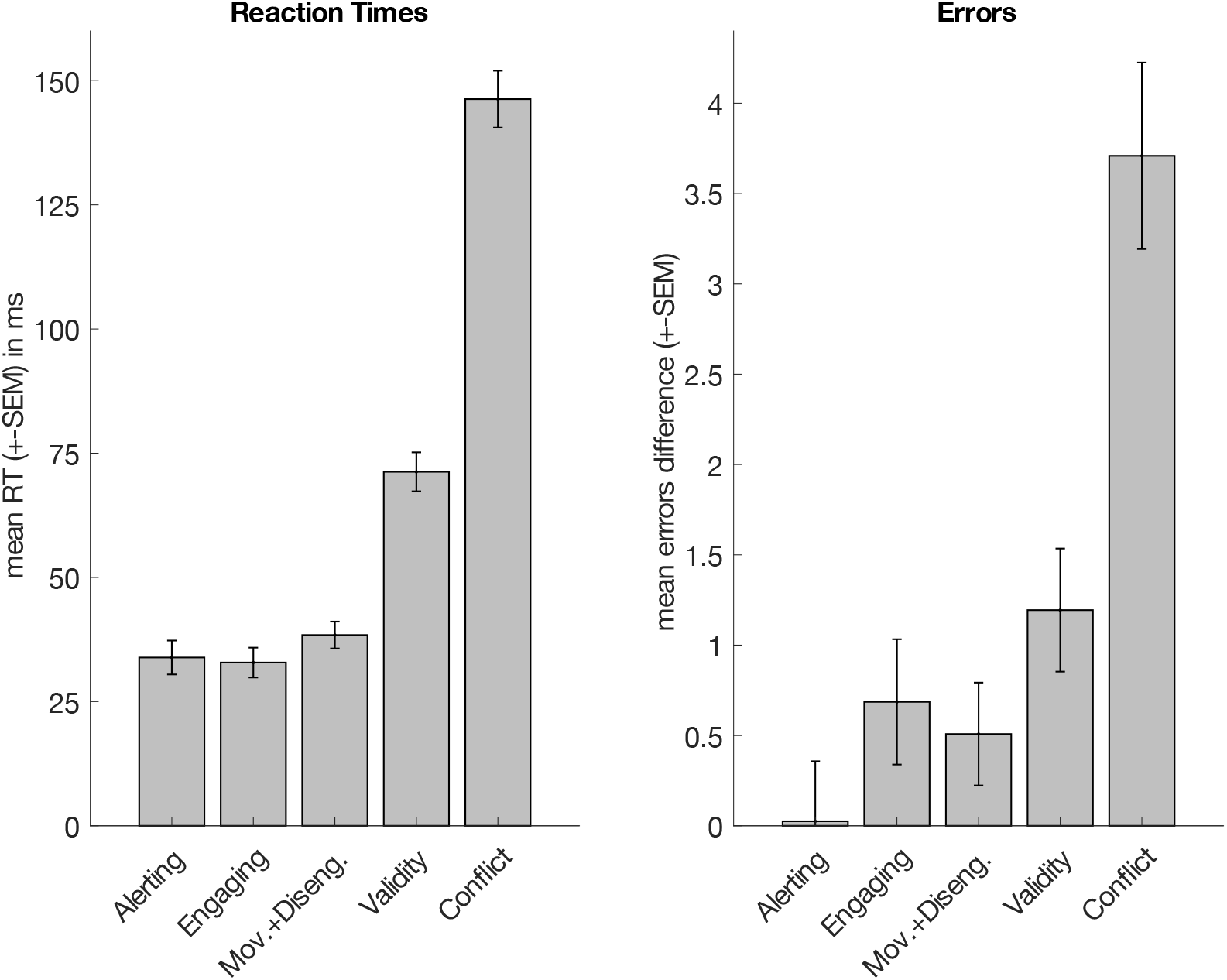
Behavioral results from the ANT-R. The bars reflect the mean difference (+SEM) between the two conditions of interest.

### 3.2 Intrinsic Connectivity Networks

Figure 3 shows the 23 group-level ICN and the hierarchical clustering of their time courses. High resolution maps of the 23 ICN are published on BALSA. Pairwise comparison of each thresholded and binarized ICN with published atlases and a list of implicated cortical regions are documented in supplementary table T1 and figure S1. The springgreen cluster on the very left of Figure 3 represents the larger *somatomotor network (SMN)*: ICN #13 (“mouth” network) as well as #15 and #21 (“hand” network). ICN #23 corresponds to the auditory network. Components #7, #12, #8 in the blue cluster on the left overlap with networks labelled cingulo-opercular, ventral attention, and salience network in different published partitions. These labels refer often to the same *midcingulo-insular network* (Uddin et al., 2019). The other two components in the blue cluster (#14 and #18) include most prominently areas along the intraparietal sulcus and correspond to the *dorsal fronto-parietal “attention” network*. The four components in the yellowgreen cluster in the middle represent the larger *visual network*: IC #6 is the primary or peripheral, and #1, #17, and #19 are the secondary or central visual network. Components #2, #4, #11 in the magenta cluster on the left represent the *default mode network*. The other component in the magenta cluster (ICN #9) is left-lateralized and overlaps with the *language network* first described in Ji et al. (2019). The Power et al. (2011) parcellation interprets this component as the ventral attention network, which has been criticized and corrected by Ji et al. (2019).

**Figure 3:**
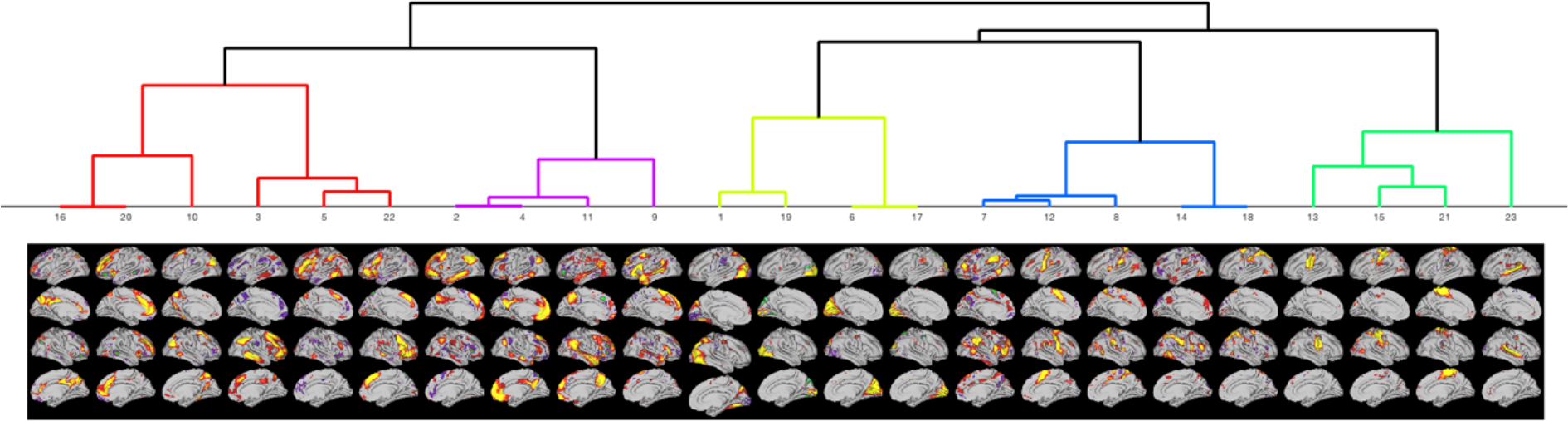
Thresholded statistical maps of independent components and their grouping into ICN through hierarchical clustering of associated time courses. The components in the red cluster belong to the executive control and fronto-parietal network, the components in magenta to the default mode and language networks, the components in light green represent the (visual) occipital network, the components in blue the midcingulo-insular and dorsal fronto-parietal “attention” networks, and the components in the darker green cluster represent the somatomotor and auditory networks. The numbers of the components correspond to the order of the ICA output (ordered by variance explained), the ordinal position in the figure was determined by the clustering.

Implicated regions are more consistent with a role in language than attention: auditory cortex, premotor area 55b, and includes inferior prefrontal cortex with areas 44 and 45 that represent Broca’s area. Components #3, #5, #22 in the red cluster on the very left correspond to the *lateral fronto-parietal network*. The remaining three components #10, #16, #20 are more difficult to label. The hierarchical clustering grouped the networks together on a hierarchy level which allowed a clear interpretation for all other components. Pairwise spatial comparisons with published templates indicate overlap with different task positive networks. Inspecting the precise boundaries of the components indicate involvement of higher order association areas including inferior and dorsolateral prefrontal cortex. We will therefore use the label *executive control network* when referring to these three components.

### 3.2 Attentional Networks

Figure 4 shows activation maps for all task contrasts (i.e. the attention networks) Alerting (a), disengaging (b), moving and engaging (c), the validity effect (d), and control (e). At large, activation patterns are consistent with the patterns reported in Xuan et al. (2016): Alerting and Control showed the most widespread activation. The different contrasts belonging to the orienting networks were more focal, which is also in line with previous reports and not surprising given that the orienting effects reflect the activation increment from spatial cues relative to the alerting effect.

**Figure 4:**
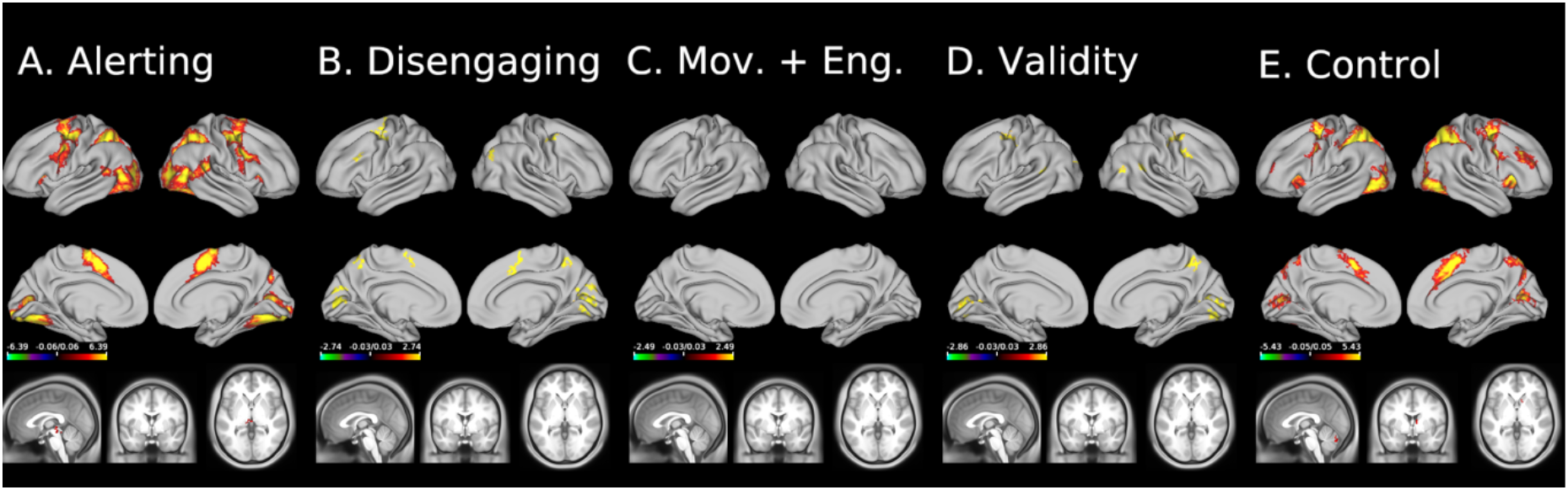
Group level activation maps (surface data) of the five contrasts. All maps show Gaussianized t-statistics, thresholded at p < .05, FWE-corrected. Underlying anatomical images are HCP’s midthickness surface mesh and HCP’s average T1-weighted structural image. We did not find significant clusters in the volumetrtic subcortical data, possibly due to the small smoothing kernel of 2 mm.

### 3.3 Spatial Regression

We fitted five linear models that regressed the spatial distribution of task-evoked activity in the five attention contrasts onto the 23 ICN. The 23 predictor variables did not show any signs of collinearity (all condition indices <1.61, all variance decomposition proportions <.5, see supplementary figure 2). The adjusted coefficients of determination (*R^2^*) of the regression models indicated that the spatial brain-wide topology of the 23 ICN accounted for 61.86% of the variance in alerting (RMSE = 1.5639,), 22.92% of the variance in disengaging (RMSE = .9897), 14.09% of the variance in moving and engaging (RMSE = .9765), 21.69% of the variance in the validity effect (RMSE = 1.0193), and 65.26% of the variance in control (RMSE = 1.1871).

Figure 5 shows regression weights for each ICN and task contrast. We will discuss involvements of ICN that explain at least 1% to the overall variance in the attention contrasts.

**Figure 5:**
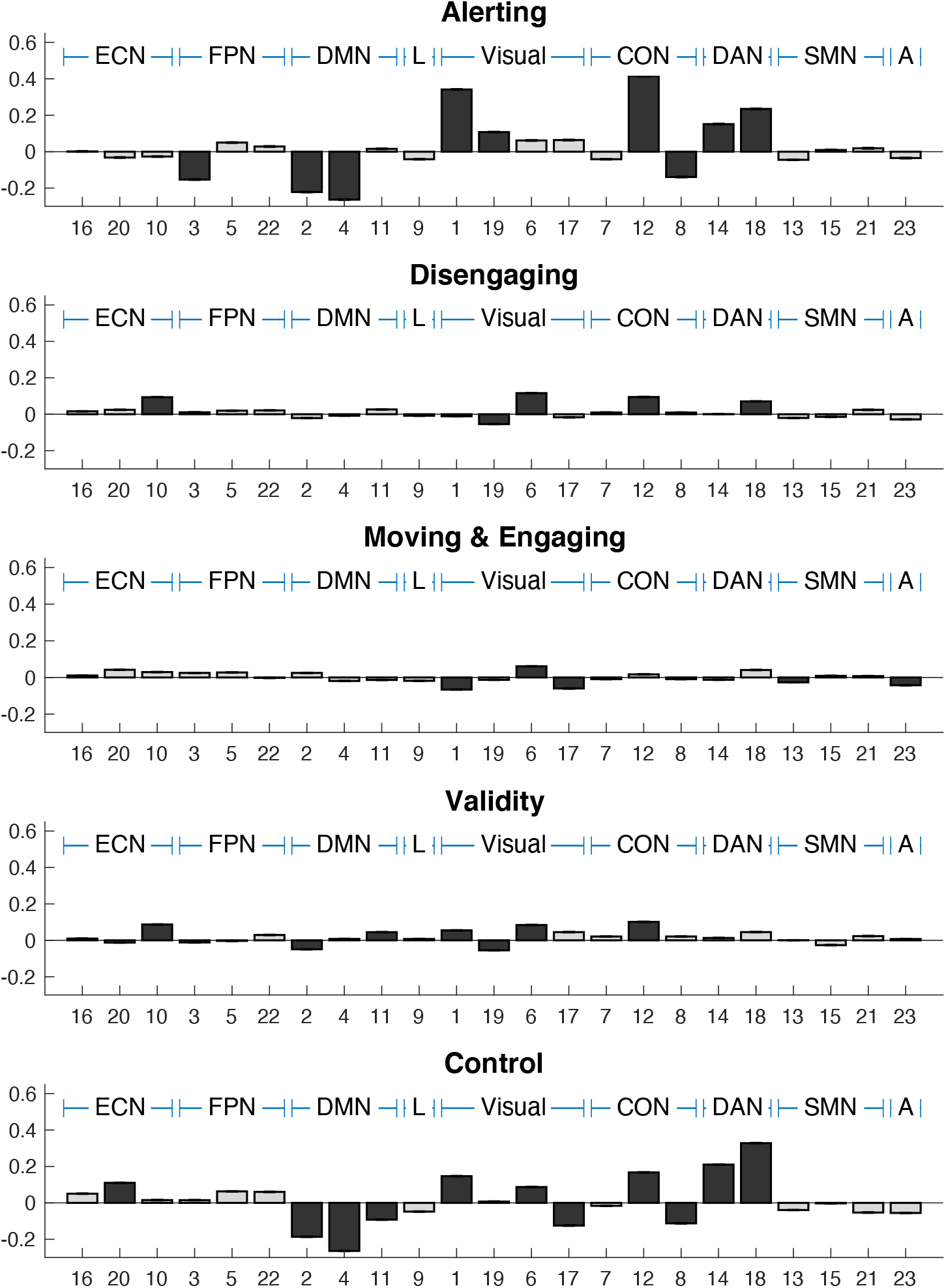
Results from the spatial regression analyses. Different panels correspond to different attention contrasts. Bars represent regression beta weights for the 23 ICN. The numbers on the x-axis reflect the order in the MELODIC. Bars in darker gray indicate components that explained at least 1% of variance in the criterion.

We first asked how many different ICN were involved in each attention contrast. Nine ICN explained at least 1% of the variance in alerting, five in disengaging, five in moving and engaging, seven in validity, and eleven in control. We then assessed whether implicated ICN were particularly enriched in any of the higher-level groups identified in the hierarchical clustering: Permutation testing revealed that the midcingulo-insular (p=.0465) and dorsal attention network (p<.001) were recruited by alerting, the visual (p=.0212) and dorsal attention network (p=.0381) by disengaging, and the visual (p<.001) and auditory network (p<.001) by moving and engaging. The visual network (p=.0034) was engaged and the default mode network suppressed (p=.0183) by the validity effect, and the visual (p<.0368) and dorsal attention network (p<.001) were engaged while the default mode network (p<.001) was suppressed by control.

We next asked whether any ICN was involved in all attention functions (alerting, control, and at least one of the orienting contrasts). We found that ICN #12 (part of the midcingulo-insular network) and ICN #18 (dorsal attention network) were recruited by all attention functions, ICN #2 (default mode network) was consistently suppressed during all attention functions, and ICN #1 (the primary visual network) was engaged by alerting, control, and the validity effect, but suppressed during moving and engaging. We then examined whether any ICN was specifically involved in only one attention contrast. This was the case for five ICN: ICN #20 (executive control network) was only activated by control, ICN #10 (executive control network) only by orienting (disengaging and the validity effect), ICN #3 (fronto-parietal network) was suppressed by alerting, and ICN #13 (“mouth” motor network) and ICN #23 (auditory network) were suppressed during moving and engaging. We also examined which ICN did not contribute to any attention contrasts: This was the case for ICN #16 (executive control network) ICN #5 and ICN #22 (fronto-parietal network), ICN #9 (language network), ICN #7 (midcingulo-insular network), and ICN #15 and ICN #21(somatomotor network).

Finally, we examined which ICN showed the largest positive contribution to each attention contrast. ICN with largest positive individual contributions were ICN #1 (primary visual network) for alerting (15.03%), ICN #6 (secondary visual network) for disengaging (6.96%), ICN #1 (secondary visual network) for moving and engaging (3.24%), ICN #12 (midcingulo-insular network) for the validity effect (3.81%), and IC #18 (dorsal attention network) for control (14.23%).

### 3.4 Overlap between attentional networks

The spatial regression analysis pointed at three ICN components that were recruited by all attention domains. Figure 6 gives a detailed picture of the ICNs and the location of activated clusters. In IC #12 and IC #18, alerting, orienting (disengaging and validity effect), and control showed overlapping activations within the superior premotor cortex: The left and right frontal eye fields (FEF) and left area 6a. The FEF and area 6a are adjacent areas, however, they belong to two different ICN: Area 6a was included in the midcingulo-insular network (IC #12) and the FEF in the dorsal attention network (IC #18). The default mode network (IC #2) was also involved in all three attention domains. Figure 5 shows that the deactivations during alerting and control followed closely the outlines of IC #2. The orienting contrasts showed less widespread deactivations at the corrected threshold which is also reflected in lower beta weights (see figure 4).

**Figure 6:**
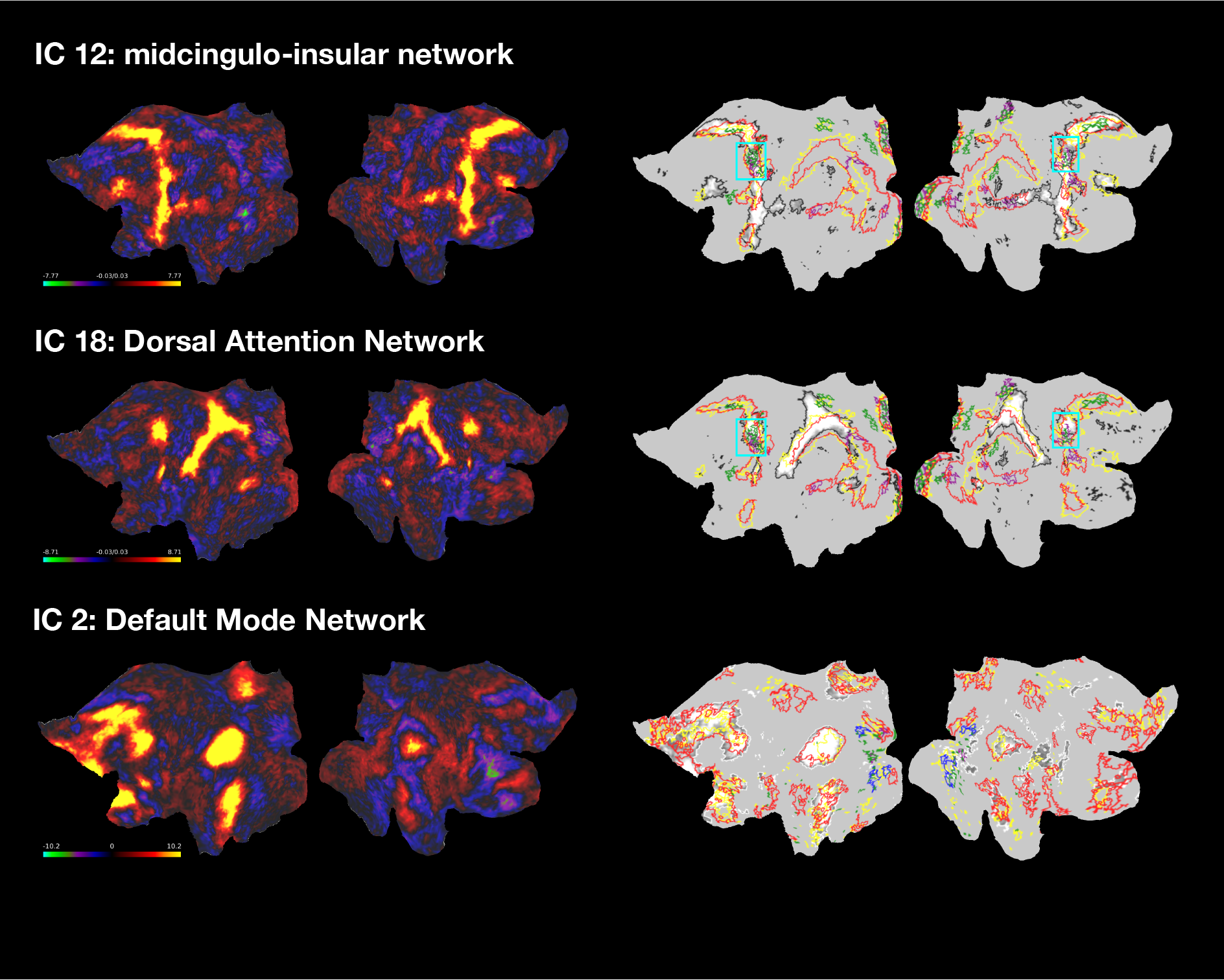
Overlapping activation in the midcingulo-insular, dorsal fronto-parietal “attention”, and default mode network. The flat maps on the left show the unthresholded IC maps. The flat maps on the right show the outline of significant task activations for alerting (red), disengaging (green), validity (purple), moving and engaging (blue), and control (yellow). Please note that the clusters in the lower panel for the default mode network correspond to deactivations. The thresholded IC maps are shown in white-gray with black outlines for spatial reference. The box in cyan highlights the frontal eye field / area 6v area where attention contrasts showed overlapping activation at the same vertex location.

## 4. Discussion

Attention network theory distinguishes three types of attention that work together in any given situation but are realized by separate networks. Different propositions have been made how these attention networks correspond to the various ICN described in the literature, yet none of them has been addressed comprehensively. To achieve a better understanding of how attention is represented at the network level, we reconstructed 23 independent components from high-resolution resting-state data and utilized a spatial regression approach (Gordon et al., 2012) to study the topological correspondence between ICNs and the activation of the attention networks. We did not find a clear correspondence between attention networks and single ICNs. Instead, we observed that each attention system activates components in multiple ICNs, that the majority (around) 70% of all components contribute to at least one attention system, and that the components activated by different attention networks overlap partially. Two components were recruited by all attention systems: A dorsal fronto-parietal network component (IC #18) which included large stretches of superior parietal as well as premotor and inferior parietal cortices and areas along the dorsal visual stream, and a midcingulo-insular component (IC #12) which included premotor and midcingular cortex, the paracentral lobule, and the posterior as well as the insular and frontal opercula. This is suggestive of a shared neural resource among the three different attention systems in the form of two separate ICN.

We grouped the 23 components into nine larger ICN and assessed whether the different attentional networks preferably recruited components from any of these ICN. We observed that the dorsal fronto-parietal network as a whole was recruited by all attention systems but that each attention system showed more widespread involvement of distinct ICN: the midcingulo-insular network in alerting, the visual and auditory networks in orienting, and visual and default mode network (deactivation) networks in control. As these findings are based on a direct comparison between the intrinsic network structure and activated attention networks, they provide important insights into the correspondence of ICN and the attention networks which may clarify previous propositions which we are going to discuss in the following.

### 4.1 Attentional networks: The extended fronto-parietal network hypothesis

It has been suggested that the fronto-parietal network underlies attention (Toro et al., 2008), which is supported by correlations between network properties of the fronto-parietal network and behavioral indices of attention (Markett et al., 2014; Visintin et al., 2015). Accordingly, Xuan et al. (2016) have argued that the three attention networks activate different parts of an extended fronto-parietal network. The fronto-parietal network described in the original reports is a larger network and hierarchically organized into separable networks: the ventral attention, dorsal attention, and fronto-parietal control network (Fox et al., 2006; Power et al., 2011; Yeo et al., 2011; Vincent et al., 2008). The ventral attention network is thought to support stimulus-driven bottom-up attention which is conceptually similar to alerting, the dorsal attention network is thought to support top-down attention which is conceptually similar to orienting, and the control network is thought to underlie executive functioning and cognitive control which is conceptually similar to the attention control system (Vincent et al., 2008; Vossel et al., 2014). While our network partition did find three larger networks that involved lateral frontal and posterior parietal cortex, they do not fully match to the three described networks.

Our dorsal fronto-parietal network (IC #14 and #18) corresponds well to the dorsal attention network. Our executive control network (IC #16, #20 and #10) includes dominantly dorsolateral and medial prefrontal cortex and the anterior cingulate and matches the description of the fronto-parietal control network. Our third fronto-parietal network (IC #3, #5, and #22), however, matches only partly the description of the ventral attention network. Our network included ventro- and dorsolateral, orbitofrontal and frontopolar cortex, as well as inferior parietal and lateral temporal cortex, but did not include the temporo-parietal junction which represents the major posterior hub in this network Fox et al., 2006; Vossel et al., 2014). In addition to the less optimal correspondence of our fronto-parietal networks with the three previously described networks, we did not observe a close fit with attention-evoked activations as well. From the three fronto-parietal networks in our partition, only the dorsal network showed clear involvement in all attention networks. Single components from the executive control network were involved in orienting (IC #10 which included mostly superior parietal and posterior cingulate cortex) and control (IC #20 which included mostly dorsolateral and anterior medial prefrontal cortex and the anterior cingulate). The third fronto-parietal network showed comparatively little involvement in any attention network, with the exception of IC #3, a right lateralized component, that was suppressed during alerting. We therefore conclude that an ‘extended fronto-parietal network’ does not capture the nature of the three attention networks well.

### 4.2 Orienting: The dorsal and ventral attention network hypothesis

The dorsal and ventral attention network have been proposed as two anatomically and functionally distinct networks (Corbetta & Shulman, 2002). In an attempt to incorporate the dorsal and ventral attention networks into attention network theory, the two networks have been equated to the orienting network (Petersen & Posner, 2012). We will first discuss the representation of the dorsal and ventral attention networks in our ICN partition before discussing their activation by the attention network test. The ventral attention network was initially described as right-lateralized but later work suggests similar organization in the left hemisphere (Vossel et al., 2014). The ventral attention network features in prominent atlases of canonical ICN (Power et al., 2011; Yeo et al., 2011) but the labeling as “ventral attention” has been contested. Others have used the labels “salience network’ (Seeley et al., 2007) or ‘cingulo-opercular network’ (Dosenbach et al., 2008) to refer to a network with similar anatomy and function. The label ventral attention network has also been used to describe a left-lateralized network whose implicated brain regions are more suggestive of an involvement in language (Ji et al., 2019; Power et al., 2011). The ventral attention network could represent a right-lateralized version of the language network (Bernard et al., 2020) but neither we nor others (Ji et al., 2019) have detected a similar right lateralized version of the language network. We decided to follow recent suggestions and use the anatomical label midcingulo-insular network (Uddin et al., 2019) for three network components that correspond closely to the cingulo-opercular network in the Cole-Anticevic and Power-partition and the ventral attention and salience network in the Yeo-partition. Of the three components, IC #7 includes bilaterally the tempero-parietal junction and ventrolateral prefrontal cortex which have been described as hubs in the ventral attention network. The identification of the dorsal attention network in our data was more straightforward. The dorsal attention network is well described in several canonical ICN atlases and we found very consistent correspondence between our components #14 and #18 and the dorsal attention network as described in these atlases.

To assess the correspondence between ICN and the orienting network, we followed previous recommendations and distinguished between different orienting effects: neural activity associated with the disengaging from an invalid spatial cue, the moving and subsequent engaging of the attentional focus to a validly cued spatial location, and the combination of the two (the validity effect) which corresponds to previously described orienting contrasts (Fan et al., 2009; Xuan et al., 2016). Our results confirm the involvement of the dorsal fronto-parietal and midcingulo-insular network in orienting. We also observed contributions from a component that we classified as part of an executive control network. This component, however, included many cortical regions that have been ascribed to the dorsal attention network in previous work (Ji et al., 2019; Yeo et al., 2011). Of note, the component within midcingulo-insular network that matched most closely the description of the ventral attention network contributed only marginally to the orienting contrasts. While the present results are well in line with the hypothesis by Petersen and Posner (2012) that the orienting network encompasses two networks that correspond to what has been described as dorsal and ventral attention network, it needs to be noted that this relationship is far from specific. We observed similar contributions of the ICN to alerting and control, which does not support the idea of a specific contribution to an anatomically distinct orienting network. Rather, our results indicate that the dorsal fronto-parietal and midcingulo-insular network play a domain-general role in the prioritization of relevant information processing that exceeds a specific contribution to the allocation of attentional resources in space.

### 4.3 Attentional Control: The fronto-parietal cingulo-opercular hypothesis

Attention network theory assumes an attention control system that is involved in the detection of targets for focal and conscious processing (Posner & Petersen, 1990), guided and controlled visual search (Posner & Dehaene, 1994), and the selection of relevant over distracting information (Fan et al., 2002). Ongoing control involves the maintenance of task-sets that set the context for moment-to-moment adjustments of cognitive processing: Previous studies indicate that set-maintenance is supported by the cingulo-opercular network and adjustments of the attentional focus is carried out by the fronto-parietal network (Dosenbach et al., 2007, 2008). This has led to the proposal that the attention control network relies on these two separate ICN: A fronto-parietal network that is distinct from the dorsal attention network and the cingulo-opercular network that we labeled midcingulo-insular network (Petersen & Posner, 2012). While our data confirm that parts of the midcingulo-insular network were recruited by the flanker contrast, we saw the strongest contributions from frontal and parietal regions that belonged to the dorsal fronto-parietal network. From the other two ICN with fronto-parietal involvement, only one component classified into the executive control network showed additional contribution. This component included dorsolateral prefrontal and anterior cingulate cortex, key regions that have been unequivocally ascribed to the attention control system (Fan et al., 2005; Fan & Posner, 2004; Petersen & Posner, 2012). While our present results are thus consistent with previous findings, they cast doubt that attentional control relies solely on a fronto-parietal network distinct from the dorsal network and the midcingulo-insular network. Rather, attentional control seems to be implemented by the dorsal attention network with additional contribution from lateral prefrontal and anterior cingulate cortex. We also observed strong deactivations of the default mode network during attentional control. The traditional view of the default mode network is that of a task-negative network that stands in an antagonistic relationship with fronto-parietal networks and de-activates unspecifically in demanding tasks (Fox et al., 2009; Raichle et al., 2001; Shulman et al., 1997). Newer evidence, however, suggests that deactivations in the default mode network encode spatial vision (Szinte & Knapen, 2020) which opens the possibility that the default mode network plays a more direct role in visual attention than expected. Future work is needed to address this hypothesis, but for now we content that the default mode network also contributes to the attentional control network.

### 4.4 Alerting and the midcingulo-insular network

Alerting refers to a state of increased sensitivity to incoming stimuli (Posner, 2008). In addition to tonic alertness as a self-initiated state of sustained vigilance, phasic alertness can use external cues to temporarily increase vigilance in anticipation of upcoming information. While both types of alertness are thought to be realized by the same alerting network, the typical alerting contrast in the attention network test uses temporal cues to induce a state of alertness and thus taps primarily into the phasic component. The midcingulo-insular network has been shown to increase its activity and functional connectivity in a task that required tonic alertness (Sadaghiani & D’Esposito, 2014) and increased pre-stimulus activity in the midcingulo-insular network leads to faster responses to unpredictable stimuli (Coste & Kleinschmidt, 2016). The midcingulo-insular network also activates in reaction to rare oddball stimuli, which implies a similar involvement in phasic alerting (Kim, 2014). We found that the midcingulo-insular network was particularly involved in the alerting contrast, supporting these previous observations. We subsumed three distinct components under the midcingulo-insular network that showed different responses in reaction to the temporal cue: The component in premotor cortex and the posterior operculum that was consistently recruited by all attention networks showed the strongest contribution to alerting. The other two components, encompassing either lateral frontal, inferior parietal, and the tempero-parietal junction or superior parietal and mid-cingulate cortices were deactivated during alerting. A suppression of activity in response to an alerting cue is usually interpreted as a preparatory reaction that facilitates faster responses (Posner, 2008) which would be a viable account for the deactivation of the two components. Additionally to the midcingulo-opercular network we found that alerting activated the dorsal attention network and deactivated a fronto-parietal component including dorsolateral prefrontal and inferior parietal cortex, as well as the default mode network. As much as we confirm the role of the midcingulo-insular network in alerting, we did not find a one-to-one correspondence between this ICN and the alerting network.

### 4.4 Overlap between attention networks

We found two ICN components that were involved in all three attention networks. While different brain regions within ICN also tend to co-activate together during tasks (Smith et al., 2009), there would still be chance that the three attention networks proposed by attention network theory dissociate within a given ICN component. But to the contrary, we found two regions in premotor cortex to be engaged by all attention systems: the bilateral frontal eye field and the left area 6v, suggesting that these two regions act as “hubs” across all attention systems. Covert spatial attention, i.e. the adjustment of the attentional focus in the absence of overt eye movements, has been tightly linked to the premotor cortex (Moore et al., 2003; Rizzolatti et al., 1987) and the frontal eye fields have been identified as the neural origin of the ‘attentional spotlight’ that modulates activity in visual areas (Thompson, 2005). Area 6v is an area in superior premotor cortex adjacent to the frontal eye field and delineates from the frontal eye field regarding its myelin content and its response profile to different tasks (Glasser et al., 2016).We found area 6v involved in the midcingulo-insular network while the frontal eye fields belonged to the dorsal fronto-parietal network. The frontal eye fields and surrounding areas have been previously associated with different attention networks (Xuan et al., 2016) but more work is needed to directly contrast the role of the frontal eye fields and area 6v within attention networks. We believe that both regions play a role in covert spatial attention by adjusting the attentional focus, irrespective of whether it is moved in space, activated in preparation of upcoming stimuli, or tuned to select relevant over irrelevant information.

### 4.5 Methodological considerations

In the following, we are going to address methodological aspects regarding the definition of attention networks, our resting-state decomposition into ICNs, and the spatial regression approach. We defined the attention networks as the set of activated grayordinates in different contrasts in the attention network test, the standard protocol proposed by the authors of attention network theory (Fan et al., 2009; Fan & Posner, 2004). This decision was motivated by previous work on attention networks (Fan et al., 2002; Xuan et al., 2016). Defining a network solely on task co-activations, however, is not without criticism. The term ‘network’ is commonly used to describe a set of network nodes including their mutual relationships (Albert & Barabási, 2002). By simply focusing on task-evoked activations we thus omitted any information on functional interactions within the attention network. Work on task-evoked whole-brain functional connectivity changes suggests that task-activations and task-connectivity carry different information (Gerchen & Kirsch, 2017) and task-connectivity can point towards important network nodes that do not show strong activation changes between task conditions (Markett et al., 2020). While the current operationalization of attention networks is thus consistent with previous work and aids interpretability in the context of previous findings, future work will want to utilize methods that aim at functional connectivity to map attention networks in more detail. When combined with analytic approaches from network science, such approach can also highlight different roles of brain area in the context of distributed systems (Zink et al., 2021).

Our main focus was a detailed comparison between activation maps from different task contrasts and the topology of ICN. While the general pattern of ICN has been well-replicated across studies, acquisition protocols, and analytical approaches, it needs to be noted that ICN are statistical abstractions of BOLD fluctuations and the exact number and topology of reconstructed ICN depends on hyperparameter choices and preprocessing strategies. Different modular decompositions of resting-state time series have been proposed throughout the literature that feature for instance six (Dosenbach et al., 2010), ten (Smith et al., 2009), seven or seventeen (Yeo et al., 2011), twelve (Ji et al., 2019), or thirteen (Power et al., 2011) ICN. In the absence of a universal ground truth, we decided to achieve our own ICN decomposition of the same participants’ resting state timeseries. We applied independent component analysis, an established approach that results in consistent and stable ICN maps (Beckmann et al., 2005), does not depend on a priori node definitions (Smith et al., 2011), allows for an automated optimization of the model order parameter (Beckmann & Smith, 2004), and most importantly, operates on the grayordinate-level which makes a direct comparison of ICN maps and task-activation maps straightforward.

Despite all progress in network neuroscience, the field has yet to agree on a comprehensive list of ICN and their names (Uddin et al., 2019). To a certain extent, the apparent differences between studies might arise from the rather indirect approach to neural activity inherent to functional neuroimaging, and from parameter choices for clustering and community detection. But more importantly, they can also reflect the hierarchical structure of functional interactions in the brain where larger networks delineate into several smaller networks at higher resolution levels (Betzel et al., 2013; Hilgetag & Goulas, 2020; Meunier et al., 2010). The hierarchical nature of ICN was also reflected in our present ICN partition. We found our reconstructed 23 signal components to correspond to nine larger ICN that have all been described in the literature. Importantly, we observed clear sensory (auditory and visual) and motor networks, which is an essential criterion for a valid network parcellation. Independent component analyses allow the different components to overlap We probed the relationship between ICNs and the attention network maps through a spatial regression approach as described previously (Gordon et al., 2012). Since we were interested in the spatial covariation of signals across the entire brain, which is expressed in single statistical parameters, no adjustment of the task activation and IC maps for multiple comparison was required and we submitted unthresholded maps to the regression analyses. We verified the absence of multicollinearity between IC components. It needs to be noted, however, that the present approach assumes static ICNs that persist across task and resting states and are invariant across participants. While these assumptions hold at large (Cole et al., 2014; Smith et al., 2009), there is still ample evidence for subtle yet reliable variation in network structure across tasks, time, and individuals (Cole et al., 2014; Feldt Muldoon, 2015; Seitzman et al., 2019). We hope that the present comparison between attention networks and the intrinsic network architecture will stipulate more research into the network-level representation of attention that will extend the current focus to temporal dynamics and individual differences.

### 4.6 Conclusions regarding attention network theory

While we found a good overall correspondence between the attention network maps and the brain’s intrinsic connectivity architecture, we did not find unique relationships between any attention network maps and single ICN, challenging most previous conjectures on the representation of attention at the network level. Each attention contrasts activated several ICN, and we found that all attention networks converged within the dorsal fronto-parietal and midcingulo-opercular network, pointing towards a shared neural resource between the different attention networks. Given that interactions and spatial overlap between attention networks have been described previously (Xuan et al., 2016), we argue to reconsider the notion of separable and independent attention networks. Instead, we propose that attention is supported by a distributed network in which different subroutines of attention (alerting, orienting, control) segregate into different subnetworks and are integrated by hubs in the dorsal fronto-parietal and midcingulo-insular network. While this proposal requires further empirical investigations, it would be well in line with several discoveries regarding the network-level representation of cognitive control and higher cognition (Braun et al., 2015; Cohen et al., 2014; Cohen & D’Esposito, 2016; Cole et al., 2013; Zink et al., 2021).

## 5. Acknowledgement

This work was funded by a grant from the German Research Foundation (DFG) awarded to Sebastian Markett (MA-6792/3-1).

