## SupplementaryMaterial for "Attention Networks and the Intrinsic Network Structure of the Human Brain"

### T1. Comparison of ICA components and resting-state networks from other parcellations.

We compared each IC component to existing network parcellations by binarizing the map at z>|3| and computing the spatial overlap with existing templates via the Jakkard index (Steen et al., 2011). Comparison with the cortical and subcortical Cole-Antcevic parcellation were done at the grayordinate-level. For the other three atlases, we converted cifti dscalar to giftis prior to comparison. The table gives the template network with the highest spatial overlap. Boldface indicates a consistent labeling across multiple parcellations.

Table T1 Spatial comparison between our ICA decomposition and existing network atlases.

| | | Cole-Anticevic | Yeo 7 | Yeo 17 | Power |
|---|---|---|---|---|---|
| red | ICN #16 | Frontoparietal | Default | Default A | Default |
| | ICN #20 | Cingulo-Opercular | FrontoParietal | SalVentAttnB | Default |
| | ICN #10 | DorsalAttn | DorsalAttn | Default C | Default |
| | ICN #3 | **Frontoparietal** | **FrontoParietal** | **Control B** | Default |
| | ICN #5 | **Frontoparietal** | **FrontoParietal** | **Control A** | **FrontoParietal** |
| | ICN #22 | **Frontoparietal** | **FrontoParietal** | **Control B** | **FrontoParietal** |
| magenta | ICN #2 | **Default** | **Default** | **Default B** | **Default** |
| | ICN #4 | **Default** | **Default** | **Default A** | **Default** |
| | ICN #11 | **Default** | **Default** | **Default B** | **Default** |
| | ICN #9 | **Language** | Default | Default B | VentralAttn |
| light green | ICN #1 | **Visual2** | **Visual** | **VisualCent** | **Visual** |
| | ICN #19 | **Visual2** | **Visual** | **VisualCent** | **Visual** |
| | ICN #6 | **Visual1** | **Visual** | **VisualPeri** | **Visual** |
| | ICN #17 | **Visual2** | **Visual** | **VisualCent** | **Visual** |
| blue | ICN #7 | Cingulo-Opercular | FrontoParietal | SalVentAttnB | FrontoParietal |
| | ICN #12 | Cingulo-Opercular | VentralAttn | SalVentAttnA | CinguloOpercular |
| | ICN #8 | SM | VentralAttn | SalVentAttnA | SM "Hand" |
| | ICN #14 | **DorsalAttn** | **DorsalAttn** | Control A | FrontoParietal |
| | ICN #18 | **DorsalAttn** | **DorsalAttn** | **DorsAttnB** | **DorsalAttn** |
| green | ICN #13 | **SM** | **SM** | **SM B** | **SM "Mouth"** |
| | ICN #15 | **SM** | **SM** | **SM A** | **SM "Hand"** |
| | ICN #21 | **SM** | **SM** | **SM A** | **SM "Hand"** |
| | ICN #23 | **Auditory** | SM | SM B | **Auditory** |

### S2. Comparison of IC components and the Glasser multimodal parcellation

We used the Glasser cortical parcellation and the corresponding mapping of the 360 cortical regions to 22 cortices to annotate the 23 ICN. The 23 ICN maps and the Glasser parcellation were converted to gifti files (one per hemisphere). The ICN maps were thresholded at Z>|4|. We then calculated for each cortex (e.g. early visual cortex) the percentage of cortical parcels that overlapped with each thresholded ICN. Results are plotted in figure S1.

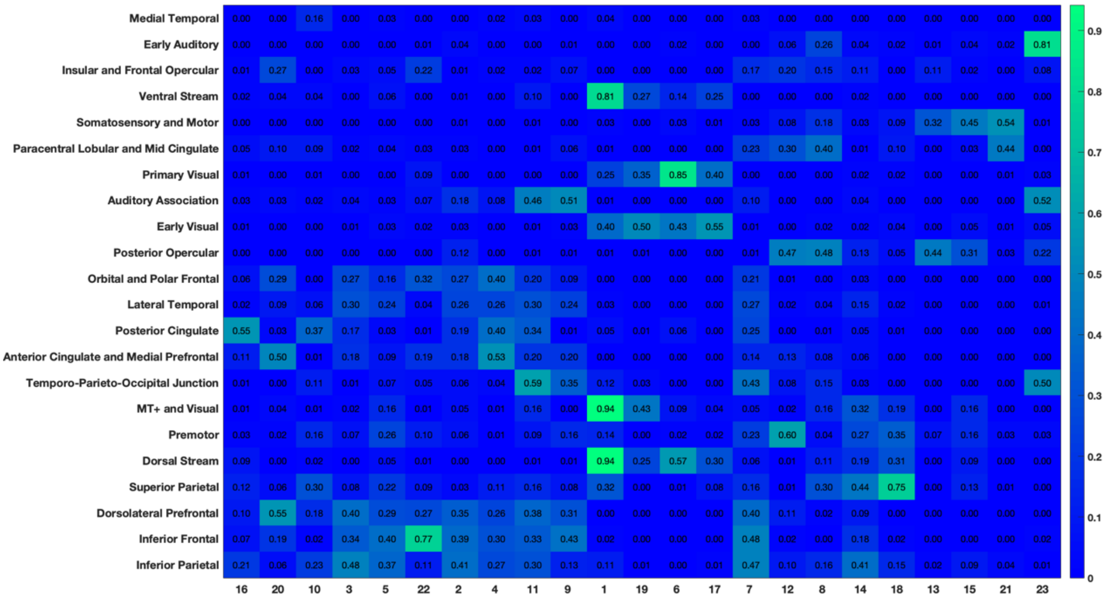


Figure S1: Overlap between ICN components and the 22 cortices (groups of cortical areas) from the multimodal parcellation.

### S2: Collinearity diagnostics

We present a tableplot (Friendly et al., 2009) with condition indices and variance decomposition proportions for each regressor (i.e. ICA component): Condition indices (the square root of the ratio of the maximum eigenvalue to each eigenvalue from the correlation matrix between standardized predictor variables) is given in the first column. By convention, condition indices <5 are considered unproblematic. The other columns give variance decomposition proportions that show the contribution of each predictor to potential variance inflation. Convention regards a predictor with two or more variance proportions >.5 as collinear. For sake of simplicity, all predictors are labeled X1, …, XN. Neither collinearity diagnostic indicated collinear relationships among predictors.


Friendly, M., & Kwan, E. (2009). Where's Waldo? Visualizing Collinearity Diagnostics. *The American Statistician, 63(1),* 56–65. https://doi.org/10.1198/tast.2009.0012

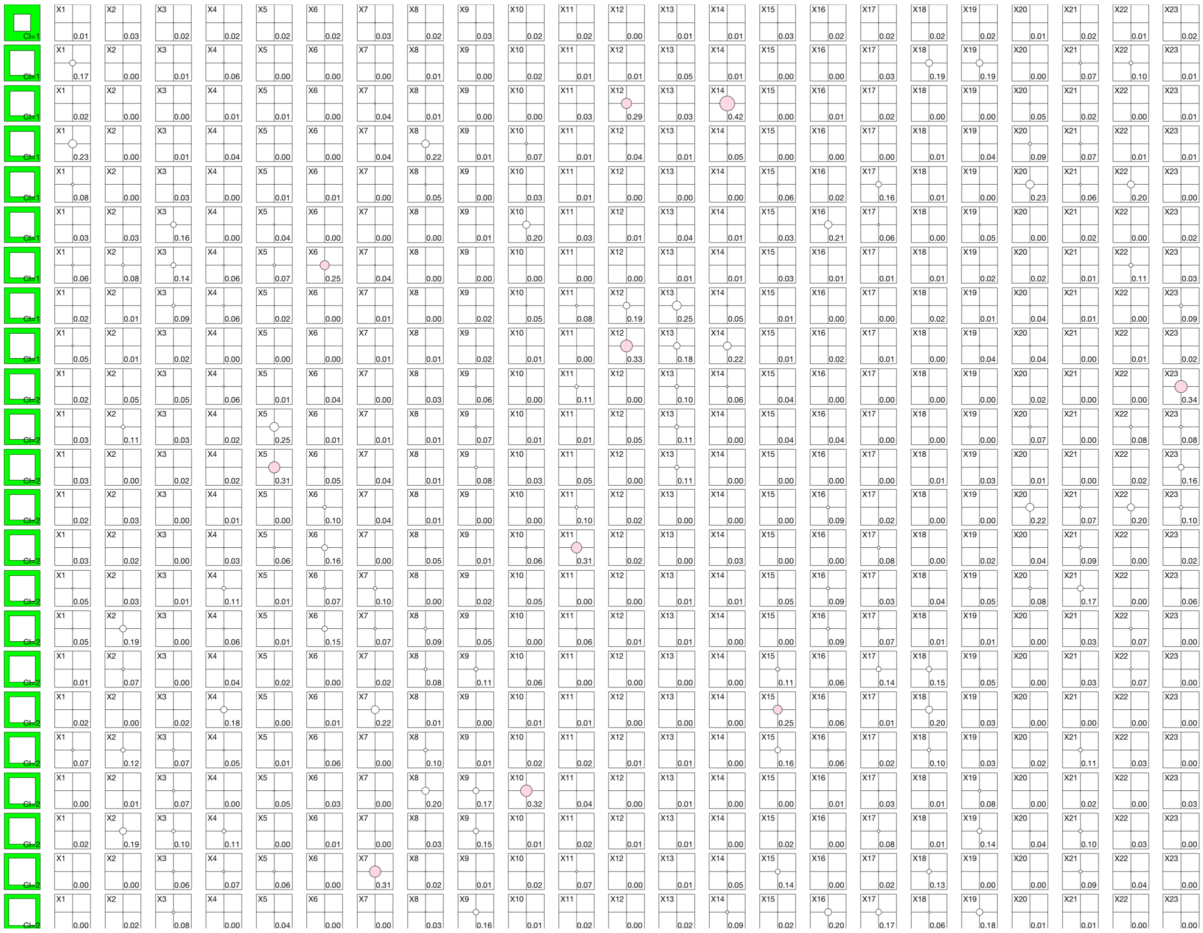
Figure S2: Collinearity Diagnostics for 23 predictors (ICA components).

### S3-S6. Regression diagnostics

We present diagnostics for the spatial regression models from the main text. We present a plot of fitted vs. actual values, a plot of the fitted values vs. the residuals, a histogram of the residuals, a plot of the residuals vs. lagged residuals (lag 1), and a probability plot for the residuals regarding normality. None of the diagnostics indicated problems for the regression analyses.

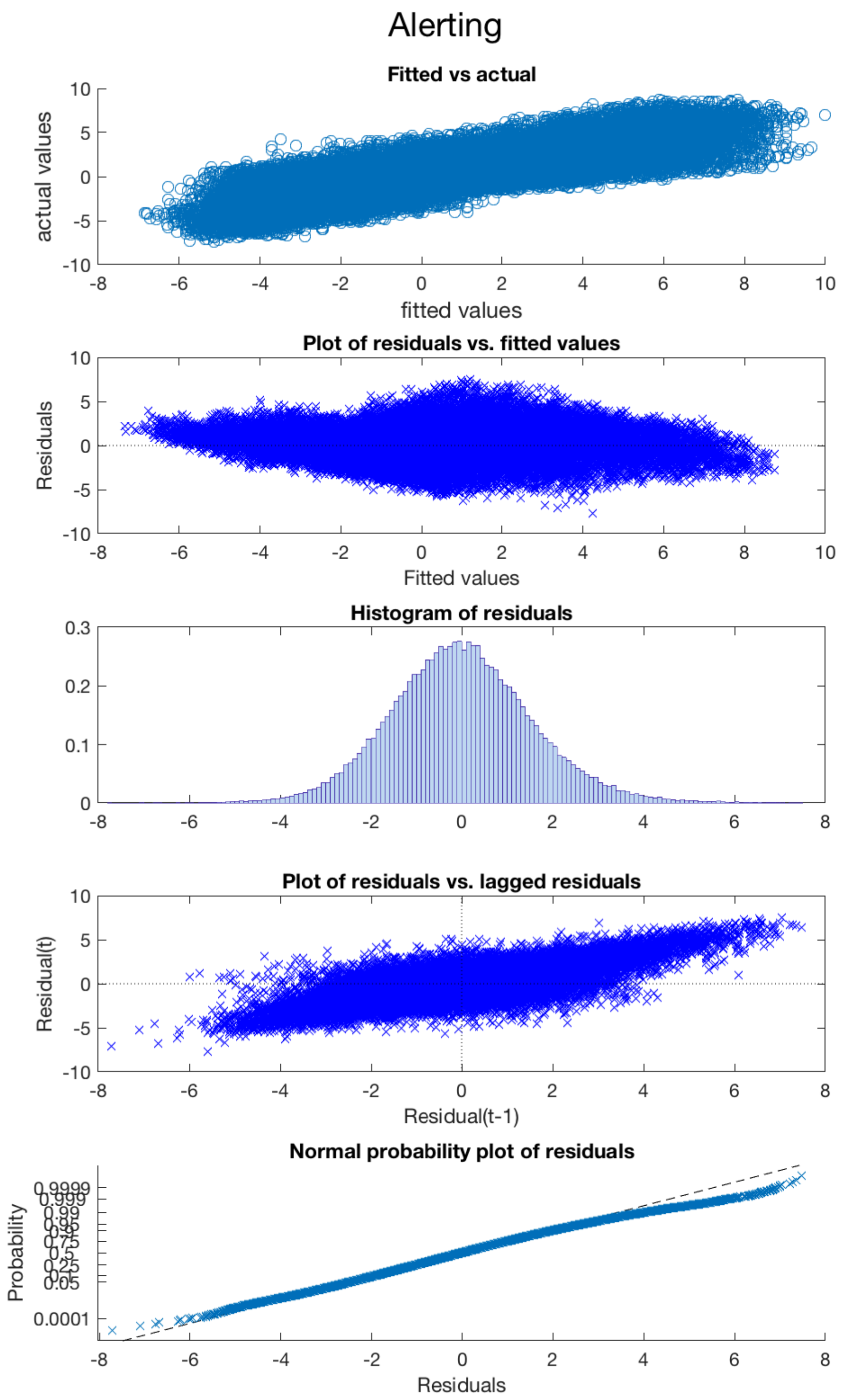


Figure S3: Regression diagnostics for Alerting

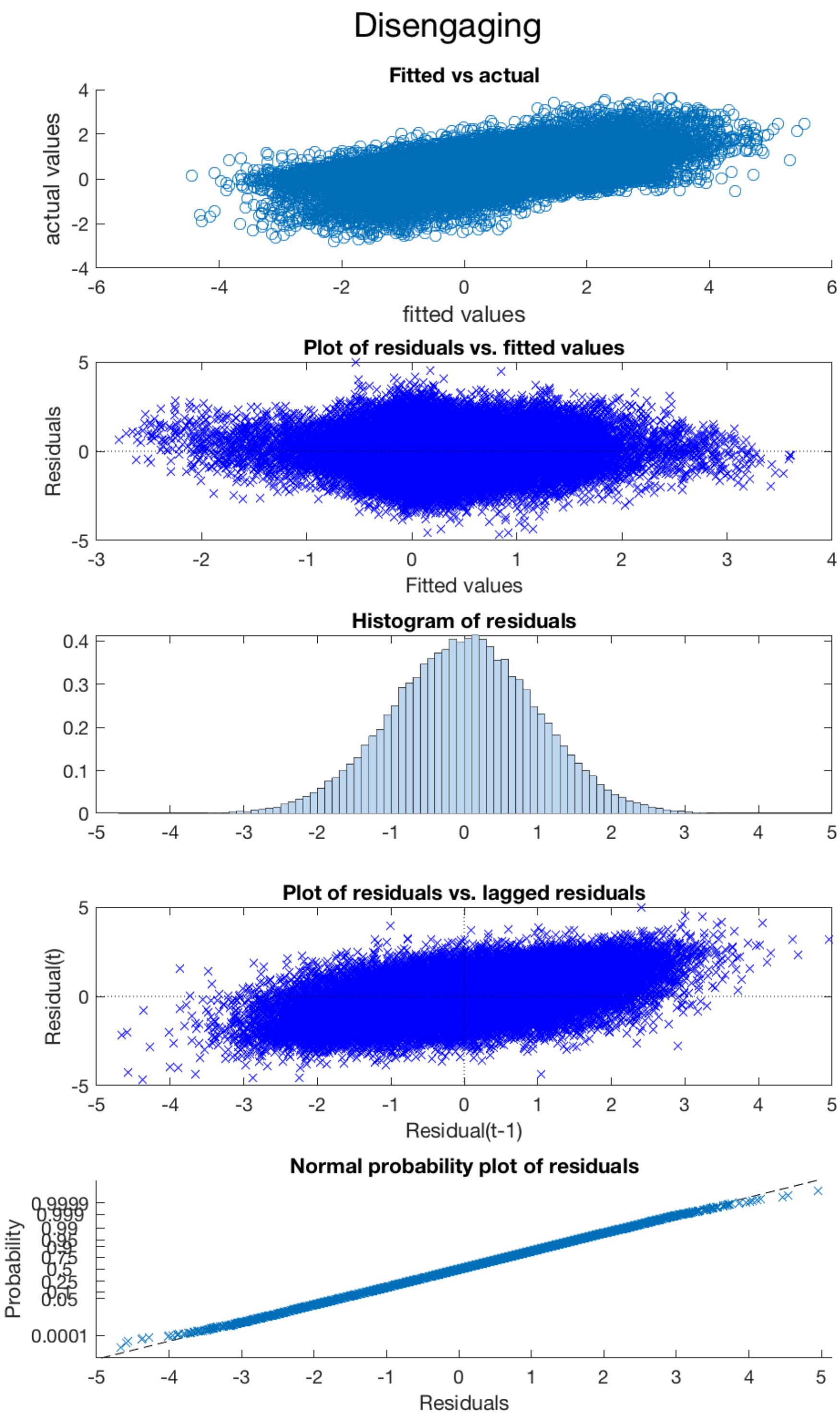


Figure S4: Regression diagnostics for Disengaging

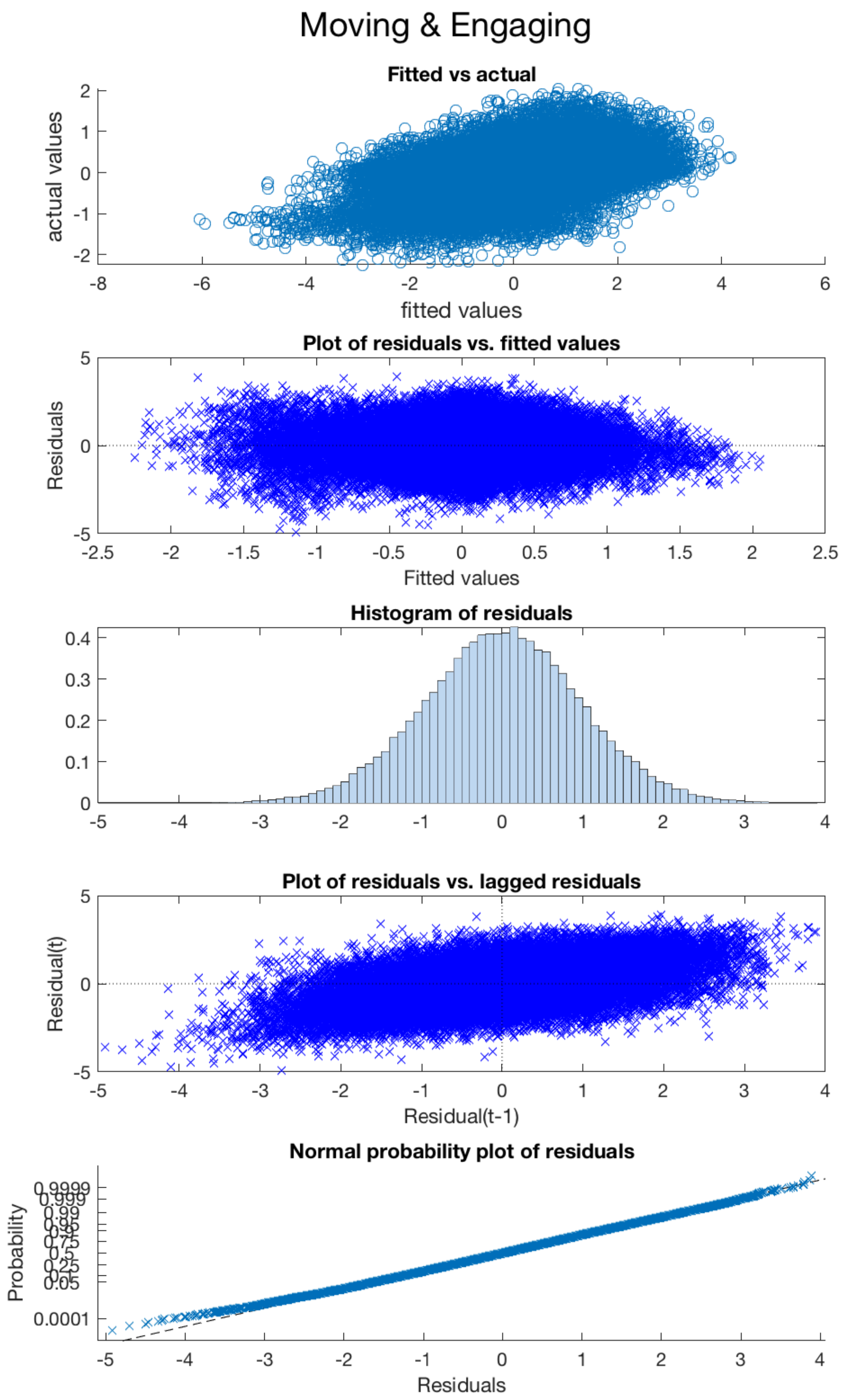


Figure S5: Regression diagnostics for Moving and Engaging

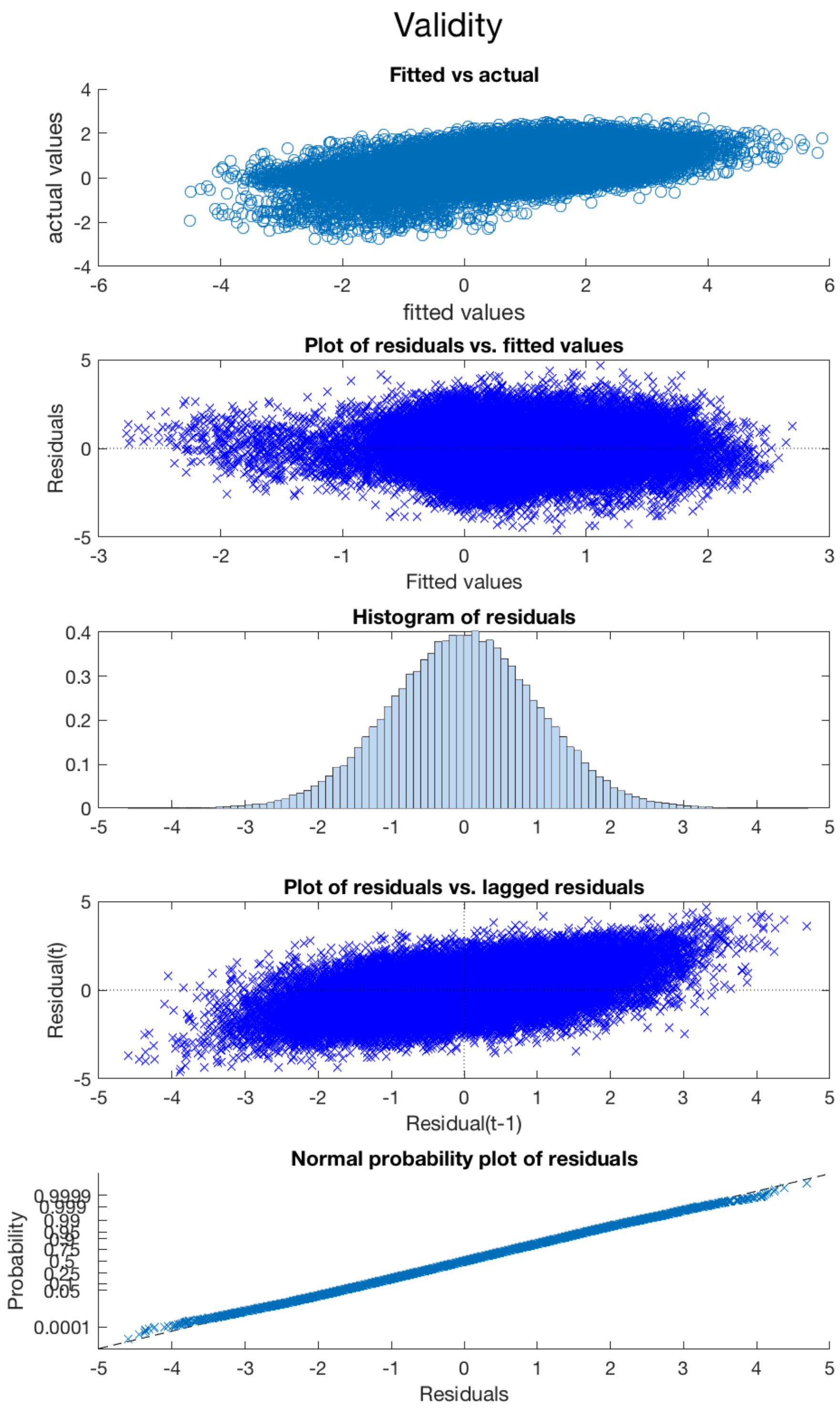


Figure S6: Regression diagnostics for the Validity effect

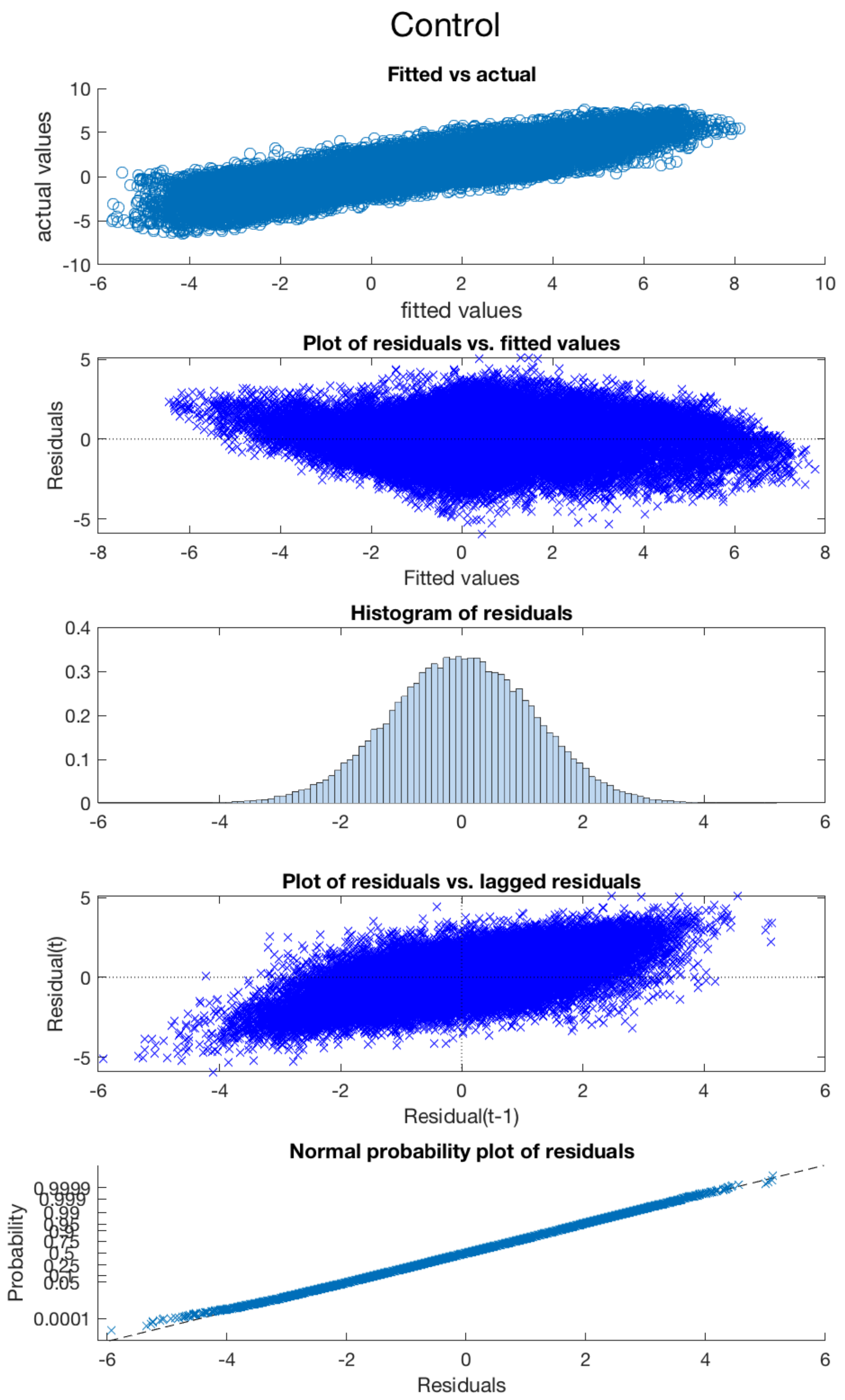


Figure S7: Regression diagnostics for control
